# Grapevine leaf MALDI-MS imaging reveals the localisation of a putatively identified sucrose metabolite associated to *Plasmopara viticola* development

**DOI:** 10.1101/2022.08.01.502001

**Authors:** Marisa Maia, Andréa McCann, Cédric Malherbe, Johann Far, Jorge Cunha, José Eiras-Dias, Carlos Cordeiro, Gauthier Eppe, Loïc Quinton, Andreia Figueiredo, Edwin De Pauw, Marta Sousa Silva

**Author notes:** Corresponding authors: M. Maia; and M. Sousa Silva, (M.S.S.). These authors are co-senior authors in this work (Edwin De Pauw, Marta Sousa Silva).

## Abstract

Despite well-established pathways and metabolites involved in *grapevine-Plasmopara viticola* interaction, information on the molecules involved in the first moments of pathogen contact with the leaf surface and their specific location is still missing. To understand and localise these molecules, we analysed grapevine leaf discs infected with *P. viticola* with MSI. Plant material preparation was optimised, and different matrices and solvents were tested. Our data shows that trichomes hamper matrix deposition and the ion signal. Results show that putatively identified sucrose presents a higher accumulation and a non-homogeneous distribution in the infected leaf discs in comparison with the controls. This accumulation was mainly on the veins, leading to the hypothesis that sucrose metabolism is being manipulated by the development structures of *P. viticola*. Up to our knowledge this is the first time that the localisation of a putatively identified sucrose metabolite was shown to be associated to *P. viticola* infection sites.

## 1. Introduction

The development of matrix-assisted laser desorption-ionisation (MALDI) mass spectrometry imaging (MSI) was first reported in 1994 and has been applied to visualise different biomolecules, since 1997 (Spengler et al., 1994; Caprioli et al., 1997). MALDI-MSI has the unique ability to analyse the sample surface directly by combining powerful raster-scan of the sample surface with lasers shots with high mass resolution mass spectrometry (Grassl et al., 2011; Bjarnholt et al., 2014; Boughton et al., 2016). This technique has not only the ability to identify specific molecules, but also to reveal the distribution of a wide range of biological compounds across the sample section in a label-free and non-targeted mode. Considering that the complexity of biochemical processes occurring in cells, tissues, organs and whole systems is not only determined by their timing, but also by the localisation of certain molecular events (Kaspar et al., 2011), MALDI-MSI has the advantage to provide high spatial resolution (Grassl et al., 2011; Kaspar et al., 2011; Bjarnholt et al., 2014; Boughton et al., 2016; Schulz et al., 2019). Moreover, MALDI-MSI is a very sensitive technique with the ability to analyse complex samples (Laugesen and Roepstorff, 2003a). The analysis of compounds in MALDI-MSI ranges from large biomolecules, such as proteins and peptides, to small molecule compounds such as lipids, sugars, amino acids, phosphorylated compounds and pharmacological and chemical compounds (Puolitaival et al., 2008; Francese et al., 2009; Goodwin et al., 2010; Solon et al., 2010; Carter et al., 2011; Goodwin et al., 2011b, 2011a; Grassl et al., 2011; Kaspar et al., 2011; Gorzolka et al., 2014a; Takahashi et al., 2015; Sarabia et al., 2018a; Alcantara et al., 2020a). MALDI-MSI is also a suitable analytical technique for both polar and nonpolar biomolecules (Rubakhin and Sweedler, 2010b).

To date, the literature of MALDI-MSI in plants is limited when compared to animal studies (Bjarnholt et al., 2014; Boughton et al., 2016). MALDI-MSI has the potential to bring new insights into the molecular analysis of plants by providing high spatial resolution information about metabolic processes and potentially determine changes during plant development or induced by environmental variation. It will provide a means of identifying the localisation of metabolites associated with tissue types, development, disease, genetic variations or following genetic manipulation (Bjarnholt et al., 2014; Boughton et al., 2016). Some reports have used MALDI-MSI to evaluate the surface distribution of sugars, metabolites and lipids in different plant tissues and organs (Bunch et al., 2004; Burrell et al., 2007; Ng et al., 2007; Li et al., 2008; Shroff et al., 2008a; Goto-Inoue et al., 2010). For instance, the distribution and profiles of metabolites were studied in barley seeds and roots to investigate the compounds associated to seed germination (Gorzolka et al., 2014b). This technique has also been used to highlight the uneven distribution of specific biological compounds in wheat grains at various developmental stages (Veličković et al., 2014). Moreover, different studies have also used this technique to better understand plant biotic and abiotic stresses (Shroff et al., 2008a; Hamm et al., 2010; Shroff et al., 2015; Soares et al., 2015; Sarabia et al., 2018b). As an example, MALDI-MSI was used in barley roots to uncover metabolites in response to salinity stress (Sarabia et al., 2018b). Also, with the increasing demands for a more sustainable agriculture practice, this technique has been used to identify specific agrochemical compounds that are present in the plant or in specific organs of the plant (Alcantara et al., 2020b).

In every mass spectrometry imaging-based analysis, the measurement of an analyte intensity is influenced by several factors including analyte extraction efficiency, ionisation efficiency and consistency of co-crystallisation with the MALDI matrix (Schwartz et al., 2003a; Schulz et al., 2019). Sample handling and preparation are crucial to obtain high quality MALDI mass spectra in a reproducible manner (Laugesen and Roepstorff, 2003a; Schwartz et al., 2003a; Grassl et al., 2011). However, the selection of the MALDI matrix and the optimisation of the preparation protocol are still empirical procedures. The composition of the solvent in which the MALDI matrix is applied on the sample influences the desorption-ionisation of molecules from the tissues. Also, the selection of the optimal matrix, including its crystallisation parameters, for the experiment depends on the type of biological molecules to be analysed. A homogeneous layer of matrix solution, followed by a fast drying of the matrix film should be preferred to avoid the formation of large matrix crystals, that are responsible for critical signal fluctuation in MALDI-MSI profile (known as the matrix effects) (Francese et al., 2009; Rubakhin and Sweedler, 2010b; Grassl et al., 2011; Kaspar et al., 2011). By its ultra-high mass resolving power, Fourier Transform Ion Cyclotron Resonance mass spectrometry (FT-ICR-MS) imaging analysis enables small molecule separation from the complex background of tissue constituents and matrix ions (Bjarnholt et al., 2014; Boughton et al., 2016), which is a very important asset when analysing complex samples such as plant tissues.

*Vitis vinifera* L. (grapevine) is one of the most important and cultivated fruit plants in the world with a highly economic impact in several countries. Unfortunately, the domesticated *V. vinifera* cultivars frequently used for wine production are highly susceptible to fungal diseases, being the downy mildew, caused by the biotrophic oomycete *Plasmopara viticola* (Berk. et Curt.) Berl.et de Toni, one of the most destructive vineyard diseases. *Plasmopara viticola* is an obligatory biotrophic pathogen. It feeds on the living tissue (grapevine) and develops specific structures to invade the cell and to obtain metabolism products, without killing the plant (Gessler et al., 2011). So far, several potential metabolic biomarkers were identified by comparing the constitutive accumulation of specific metabolites not only in the leaf tissue of different *Vitis* genotypes (Maia et al., 2020), but also in grapevine-pathogen interaction (Batovska et al., 2008, 2009; Becker et al., 2013; Adrian et al., 2017; Nascimento et al., 2019). Their accumulation reflects however the all-leaf tissue metabolome composition/modulation and not the molecules that contribute for the first contact with pathogens at leaf surface. Some studies in grapevine leaves with MSI are starting to appear to identify the main biological compounds in the leaf surface during the interaction with *Plamopara viticola* (Hamm et al., 2010; Becker et al., 2014, 2017). However, these studies are only focused on the detection of specific compounds already described to have a defence role on grapevine after pathogen infection, e.g., resveratrol, pterostilbene and viniferins. Also, previous studies have reported that the lower side surface of leaves of *Vitis* genotypes differ distinctively with respect to trichomes. The distribution of these structures appears to be very characteristic of each genotype ranging from complete absence to dense coverage (Konlechner and Sauer, 2016).

The present work aimed at analysing *V. vinifera* ‘Trincadeira’ leaf surface with and without *P. viticola* infection, using MALDI FT-ICR-MS imaging approach. Sample preparation, including trichomes removal, the choice of the MALDI matrix and the matrix deposition solvent, were optimised to uncover the localisation of biological compounds present after pathogen infection.

## 2. Materials and methods

### 2.1. *P. viticola* propagation

*Plasmopara viticola* sporangia were collected from infected plants from the vineyard at the Portuguese Ampelographic Grapevine Collection (CAN, international code PRT051, established in 1988) at INIAV-Estação Vitivinícola Nacional (Dois Portos), using a vacuum pump (Millipore) at 10 kPa and stored at −20 °C. Inoculum was propagated in *V. vinifera* ‘Carignan’ (susceptible cultivar) leaves by spreading a sporangia solution on the lower side surface of the leaf using a sterilized laboratory spreader, with an undefined concentration of *P. viticola* sporangia. After infection, leaves were kept in the dark for the first 8 to 12 hours and then kept in a climate chamber with controlled conditions (25 °C; 16/8 hours with light/dark; 90% humidity) until sporulation appeared in all leaf surface. Sporangiophores with sporangia were collected with a vacuum pump (Millipore) at 10 kPa and stored at −20 °C until further use.

### 2.2. Plant material harvesting and *P. viticola* infection assay

The third to fifth fully expanded leaves, from shoot to apex, of *Vitis vinifera* ‘Trincadeira’ (VIVC variety number: 15685) at the veraison stage, susceptible to *P. viticola*, collected at the Portuguese Ampelographic Grapevine Collection. CAN is located at Quinta da Almoinha, 60 km north of Lisbon (9° 11’ 19” W; 39° 02’ 31” N; 75 m above sea level), and occupies nearly 2 ha of area with homogeneous modern alluvial soils (lowlands) as well as drained soil. All accessions are grafted on a unique rootstock variety (Selection Oppenheim 4–SO4) and each accession come from one unique plant collected in the field. The climate of this region is temperate with dry and mild summer. The best possible health status was guaranteed for all accessions

For leaf infections, sporangia viability was confirmed by microscopic observations as described by Kortekamp and co-workers (Kortekamp et al., 2008), prior to inoculation. One mL of a suspension containing 50 000 spores/mL was spread all-over the lower side of the leaf using a sterilized laboratory spreader. After inoculation, plants were kept for 8 to 12 hours in the dark and kept under natural light conditions at 25 °C (90% humidity) for 96 hours. Two types of inoculated leaves were obtained for analysis: leaves infected with *P. viticola* without visible sporulation and with visible sporulation. Leaf discs of 1 cm were then cut using a cork-borer and lyophilised at −50 °C between 2 glass slides, to obtain a planar surface once dried. Mock inoculations (controls) were performed by rinsing *Vitis vinifera* ‘Trincadeira’ leaves with water and keeping them for 8 to 12 hours in the dark. After that, control leaves were kept under natural light conditions at 25 °C for 96 hours. Leaf discs of 1 cm were cut using a cork-borer and lyophilised at −50 °C between 2 glass slides, to obtain a planar surface once dried. Infection control was accessed 8 days after inoculation with the appearance of typical disease symptoms (**Figure S1**).

### 2.3. Matrix coating optimisation

MALDI matrices α-cyano-4-hydroxycinnamic acid (HCCA), 2,5-dihydroxybenzoic acid (DHB) and 9-aminoacridine (9-AA) were purchased from Sigma-Aldrich (Overijse, Belgium). Trifluoracetic acid (TFA) was from Sigma-Aldrich, and LC-MS grade acetonitrile (ACN) and methanol (MeOH), were purchase from Biosolve (Valkenswaard, Netherlands). Indium tin oxide (ITO)-coated glass slides were purchased from Bruker Daltonics (Bremen, Germany). For matrix coating optimization on leaf discs with and without trichomes, lyophilised control *Vitis vinifera* ‘Trincadeira’ leaf discs were carefully removed from the plates and transferred to the target ITO-glass slide, previously covered with a double-sided adhesive copper tape (StructureProbe INC, West Chester, PA, USA), with the lower side surface of the leaf facing up. For the analysis of trichome-free leaf discs, after mounting, trichomes were carefully removed using tweezers under a magnifying lens. After trichomes removal, microscope images (obtained at the magnification 10 x with an Olympus BX40 microscope) of the leaf discs were performed to visualise the leaf discs for possible damages. Only not damaged and flat areas of the microscopic images were selected for MSI analysis. Both leaf discs were sprayed with HCCA matrix at 5 mg/mL using a solution of 70:30 (v:v) ACN:H_2_O (**Table 1 – (1)**).

**Table 1.** Different matrices, concentrations and layers used in *Vitis vinifera* ‘Trincadeira’ leaf discs.

|  | Matrix | Concentration | Solvent (% v/v) | Matrix application method | Number of layers |
| --- | --- | --- | --- | --- | --- |
| (1) | HCCA | 5 mg/mL | 70% ACN<br>30% H<sub>2</sub>O<br>0.1% TFA | SunChrom | 20 |
| (2) | HCCA | 5 mg/mL | 70% MeOH<br>30% H<sub>2</sub>O<br>0.1% TFA | SunChrom | 20 |
| (3) | DHB | 20 mg/mL | 50% ACN<br>50% H<sub>2</sub>O<br>0.1% TFA | SunChrom | 50 |
| (4) | 9-AA | 15 mg/mL | 100% MeOH<br>0.1% TFA | Commercial spray | 15 |

After optimization, *Vitis vinifera* ‘Trincadeira’ leaf discs: control (mock inoculated), 96 hpi (hours post-inoculation) with *P. viticola* without visible sporulation and 96 hpi with *P. viticola* visible sporulation were transferred to ITO-glass slides and trichomes were removed. ITO-glass slides were stored in a desiccator until matrix deposition. Leaf discs were sprayed with the different matrices in different concentrations and solvents (**Table 1**). HCCA matrices of 5 mg/mL were prepared using 70:30 (v:v) ACN:H_2_O and 70:30 (v:v) MeOH:H_2_O with 0.1% (v/v) TFA. In total, 20 layers of this matrix solution were sprayed on the ITO slide using the SunCollect instrument (SunChrom, Friedrichsdorf, Germany). The first layer was sprayed at a flow rate of 10 μL/min. Flow rate was increased by 10 μL/min after each layer until reaching 60 μL/min. DHB matrix was prepared in 50:50 (v:v) ACN:H_2_O with 0.1% (v/v) TFA to reach a final concentration of 20 mg/mL according to Seaman and co-workers (Seaman et al., 2014) and fifty matrix layers were sprayed on the ITO slide, as previously described. Solution of 9- AA was prepared at 15 mg/mL in methanol with 0.1% (v/v) TFA according to Rohit Shroff and co-workers (Shroff et al., 2008b). Leaves were spray-coated by using a commercial sprayer. The target plate was kept at around 45 ° angle and the sprayer held at 20 cm from the plate. This insured that the cone of the spray reaching the target covered the entire leaf. Each spray was followed by 10 seconds of warm air drying. This process was typically carried out 15 times to give maximal signal strength.

### 2.4. MALDI FT-ICR-MS imaging analysis

Microscope images (obtained at the magnification 10 x with an Olympus BX40 microscope) of leaf discs onto ITO-glass slides were acquired to better visualise and select areas of interest in the discs. Also, since the leaf surface is not entirely flat, only the flat areas of the microscopic images were selected for MSI analysis. Small areas of each leaf disc were selected to be analysed (**Figure S2**). Analysis was performed using a SolariX XR 9.4T FT-ICR-MS (Bruker Daltonics, Bremen, Germany), fitted with the dual ESI/MALDI ion sources and SmartBeam laser. For each mass spectrum, the following laser parameters were used: 200 laser shots at a repetition rate of 1000 Hz, 60% of power. The raster step size was set at 200 μm. Mass spectrometry images were acquired in positive ionisation mode, in the mass range of 200 to 1000 *m/z.* Red phosphorus solution in pure acetone spotted directly onto the ITO Glass slide was used to calibrate the mass spectrometer before each analysis.

### 2.5. Data analysis

Data were analysed using the MSI software SCiLS^™^ Lab 2016b (Bruker Daltonics, Bremen, Germany) for unsupervised spatial identification of discriminative features between control, 96 hpi, and 96 hpi with visible sporulation of *P. viticola* sporangiophores. For each pixel, data was normalised by the total ion count. MALDI FT-ICR-MS chemical images were generated from a colour scale, which represents the normalised intensity of specific ions. Each pixel of the image is associated with the original mass spectrum that is acquired in a particular position. A spatial localisation of the analytes is provided as a function of the *m/z* values. For metabolite identification, selected *m/z* values were submitted to MassTRIX 3 (Suhre and Schmitt-Kopplin, 2008) server (http://masstrix.org, accessed in March 2022) considering the following parameters: positive scan mode; the adducts [M+H]^+^, [M+K]^+^ and [M+Na]^+^ were considered; a maximum *m/z* deviation of 2 ppm was considered; *Vitis vinifera* was selected; search was performed in the databases “KEGG (Kyoto Encyclopaedia of Genes and Genomes) /HMDB (Human Metabolome DataBase)/LipidMaps without isotopes”.

## 3. Results

### 3.1. Preparation of grapevine leaves for analysis: the influence of trichomes

Grapevine leaves have trichomes in the lower side surface (Konlechner and Sauer, 2016), which may hamper the analysis and influence the MSI results (Bjarnholt et al., 2014). Hence, we started by analysing grapevine control leaf discs with and without trichomes to understand if these structures influence the coating of the matrix and to select the leaf discs leading to the best matrix coating. For this analysis, α-cyano-4-hydroxycinnamic acid (HCCA) was chosen as the matrix as it has been described in the literature to form small crystals, producing a more homogeneous matrix coating (Laugesen and Roepstorff, 2003b; Schwartz et al., 2003b). HCCA was prepared according to (1) in **Table 1 (Materials and Methods section**).

*V. vinifera* ‘Trincadeira’ leaf discs’ areas analysed with and without trichomes are presented in **Figure 1**. In positive-ion mass spectrum, the global intensity of the peaks detected was higher in the leaf disc without trichomes and with a more homogenous intensity detection. These results demonstrate that the matrix was better distributed in the leaf disc without trichomes and consequently a better signal was detected. For further analyses, trichomes were removed from all leaf discs using a tweezer under a magnifying lens. Moreover, to avoid leaf topological interference with the laser, only flat areas of the microscopic images (i.e., corresponding to flat regions on the microscopic image) were selected for MSI analysis.

**Figure 1.**
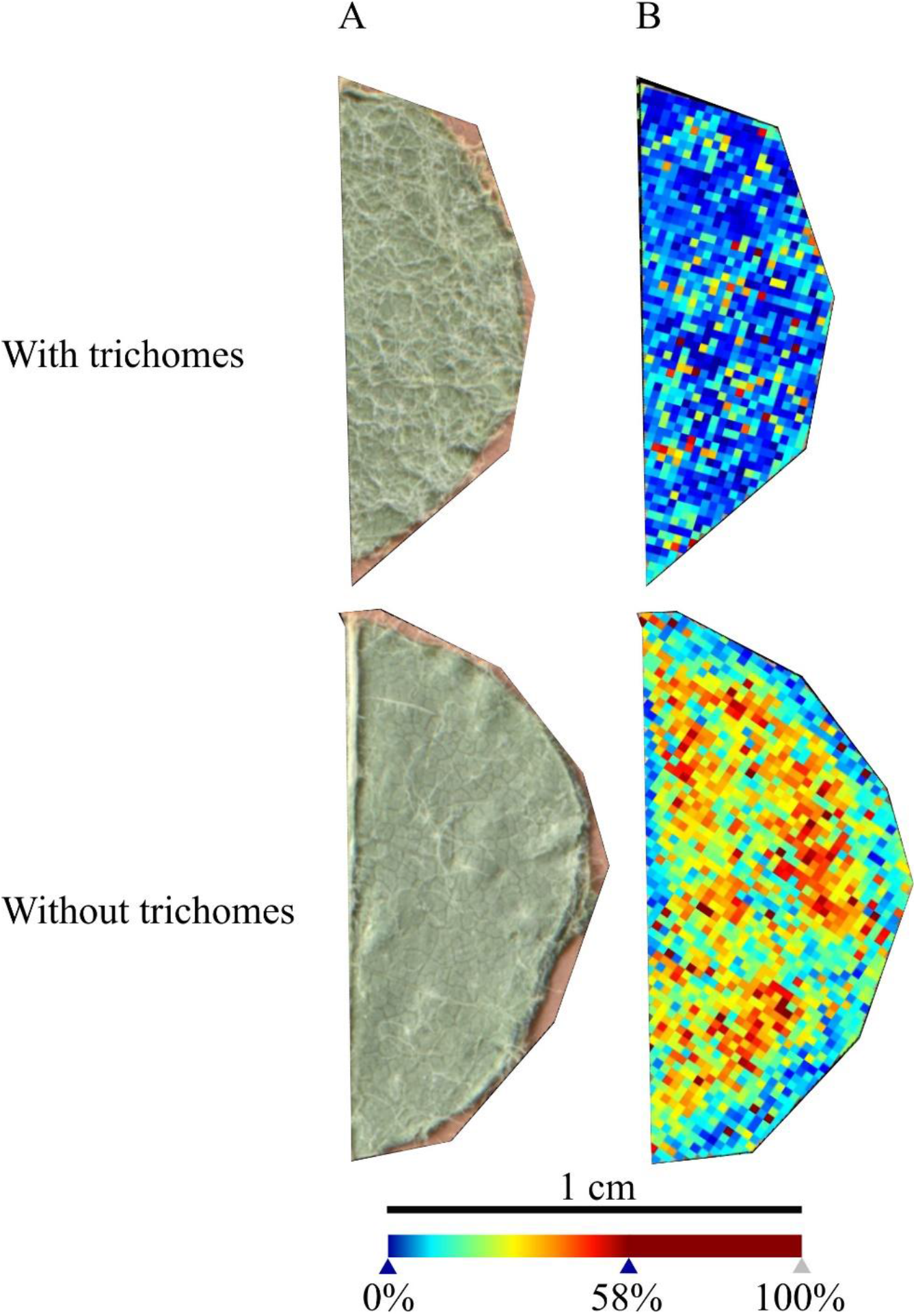
*Vitis vinifera* ‘Trincadeira’ leaf discs analysed with and without trichomes. **A,** Microscopy images of the leaf disc sections analysed. **B,** MALDI-FT-ICR-MS images reconstructed from all the peaks detected in the leaf discs. Spatial resolution: 30μm.

### 3.2. Different matrix analysis and metabolite detection

After plant material preparation optimisation, and to detect molecules related to the grapevine-*P. viticola* interaction, we tested different matrices already described for MALDI FT-ICR MS imaging analysis of plant tissues (**Table 1**).

Matrices α-cyano-4-hydroxycinnamic acid (HCCA), 2,5-dihydroxybenzoic acid (DHB) and 9- aminoacridine (9-AA) were prepared according to **Table 1** and applied by spraying on the different *V. vinifera* ‘Trincadeira’ leaf discs: control (mock inoculated), 96 hpi (hours post-inoculation) with *P. viticola* without visible sporulation and 96 hpi with *P. viticola* visible sporulation. HCCA was prepared into two different solvents: MeOH and ACN. HCCA is the most commonly used matrix for metabolites and different MALDI imaging studies in plant tissues were reported using that matrix in both solvents (Costa et al., 2013; Becker et al., 2014; Seaman et al., 2014; Soares et al., 2015). Microscope images (10x) of leaf discs were taken for a better selection of the leaf discs areas to be analysed (**Figure S2**). Selected areas were analysed with an FT-ICR-MS and ion images were reconstructed from the absolute intensities of the mean mass spectra, showing the exact location of ions on the extension of the leaf area analysed. The mass spectra of grapevine leaf discs areas obtained from HCCA matrix with MeOH and ACN as solvents were very similar (**Figure S3**). These results demonstrate that both solvents can be used for MALDI analysis of grapevine leaf discs. Comparing all spectra, the intensity of the peaks, within the 800 to 900 *m/z* range, was lower for the matrices DHB and 9-AA (**Figure S3**).

These results also confirm our previous analysis. Removing the physical barrier from the grapevine leaves, trichomes, allows for a better MSI analysis independently of the matrix and solvents used.

Reconstructed MS images by MS data of all matrices, demonstrated that the ion intensity at *m/z* 365.105 (1.1 ppm) and 381.079 (1 ppm) is higher in the leaf discs infected by *P. viticola* than in the control discs (**Figure 2**). The highest signals were observed with HCCA in MeOH and, to a lower extent with HCCA in ACN. For DHB and 9-AA, the signal was found much lower and less detectable than with the other matrices.

**Figure 2.**
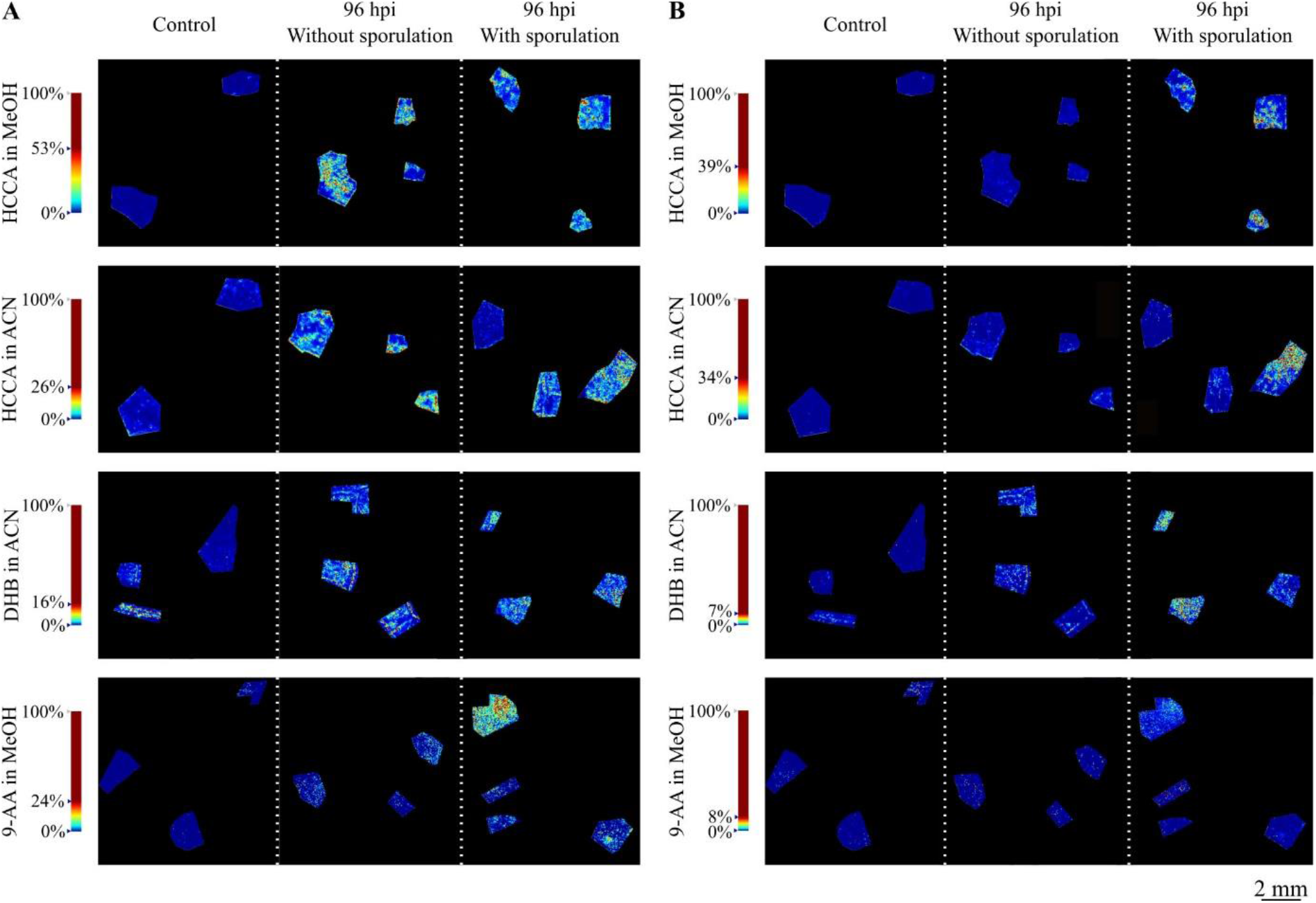
Reconstructed ion images of putatively identified sucrose. **A,** (m/z 365.105, [M+Na]+) and **B,** (m/z 381.079, [M+K]+) adducts, detected with MALDI-FT-ICR-MS imaging using HCCA matrix with MeOH and ACN, DHB matrix and 9-AA matrix. Leaf disc areas analysed from control (mock inoculated), 96 hpi without visible *P. viticola* sporulation and 96 hpi with visible *P. viticola* sporulation are presented. The colour scale of the leaf disc areas indicates the absolute intensity of each pixel (arbitrary units). Spatial resolution: 30μm.

For compound annotation, MassTRIX database was used and *m/z* 365.105 and 381.079 ions were putatively identified as the [M+Na]^+^ and [M+K]^+^ adducts, respectively, of a disaccharide with a molecular formula of C_12_H_22_O_11_. Being sucrose the most common disaccharide in plants and given our previous results (Nascimento et al., 2019) showing that sucrose increased in *Plasmopara viticola*-infected grapevine leaves 24h post-inoculation, we can attribute the ion images of *m/z* 365.105 and 381.079 to putatively identified sucrose. In a recent study based on the accurate mass measurements by Taylor and co-workers, these masses were also detected and correspond to different adducts of disaccharides (Taylor et al., 2021).

In a more detailed analysis, it was visible that the spatial distribution of putatively identified sucrose in the infected grapevine leaf discs is non-homogeneous, as observed in **Figure 2**. To evaluate this distribution, the accumulation of these ions was further investigated in the network of small veins and more precisely in the dense parenchyma tissue (**Figure 3** and **Figure S4**). Images of the 96 hpi without visible *P. viticola* sporulation and 96 hpi with visible *P. viticola* sporulation leaf discs were reconstructed from the putatively identified sucrose *m/z* 365.105 ([M+Na]^+^) as the distribution for *m/z* 381.079 ([M+K]^+^) was similar but less intense. Our results suggest that putatively identified sucrose, *m/z* 365.105 ([M+Na]^+^) and *m/z* 381.079 ([M+K]^+^), are mainly located around the veins, which is an indicator of putatively identified sucrose at *P. viticola* infection sites (**Figure 3** and **Figure S4**).

**Figure 3.**
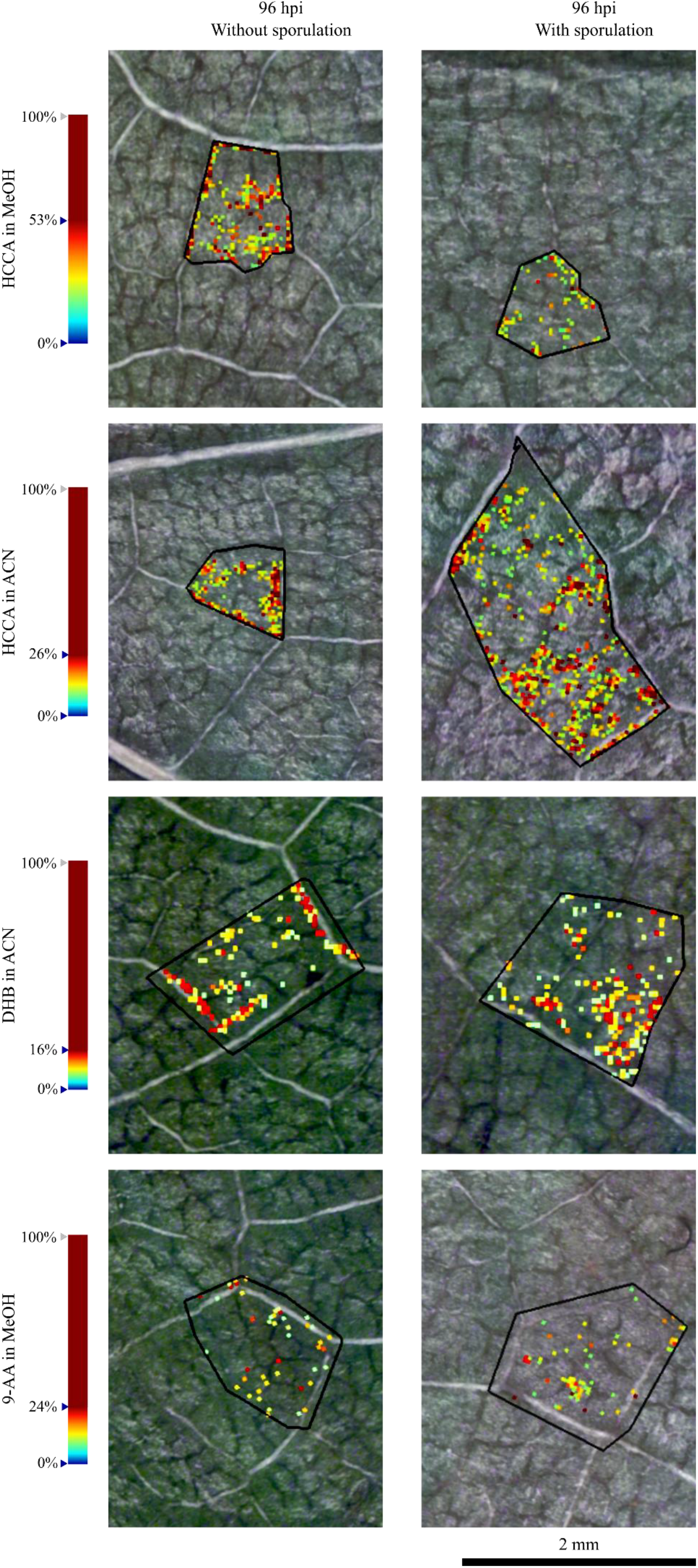
Reconstructed ion images of one selected region of putatively identified sucrose *(m/z* 365.105, [M+Na]^+^) in 96 hpi without visible *P. viticola* sporulation and 96 hpi with visible *P. viticola* sporulation. Leaf disc areas detected via MALDI–FT-ICR-MS imaging using HCCA matrix with MeOH and ACN, DHB matrix and 9-AA matrix. The colour scale indicates the absolute intensity of each pixel (arbitrary units). White arrows indicate the visible distribution of putatively identified sucrose along the leaf veins. Spatial resolution: 30μm.

## 4. Discussion

Mass spectrometry imaging has been applied to study several plant tissues. However, sample preparation of plant tissues is not easy and continuously seems to be the bottleneck of this technique (Dong et al., 2016). This topic has been the major area of interest with the goal to improve sensitivity, specificity and obtain good quality images in a reproducible manner (Schulz et al., 2019).

Sample preparation is a particular challenge, because the visualisation of certain classes of compound relies on specific conditions for optimal ionisation. Careful sample treatment is essential for signal quality and the avoidance of lateral displacement of the analytes (Rubakhin and Sweedler, 2010a). Artefacts can arise at any stage between sample collection and MSI analysis (Rubakhin and Sweedler, 2010a). Although different methodologies have been published through the years to limit these issues, the optimisation of sample preparation is still sample dependent, because of inherent differences in the composition of plant tissues (Grassl et al., 2011; Kaspar et al., 2011).

In plants, trichomes are essential epidermal outgrowths covering most aerial plant tissues and can be found in a very large number of plant species (Huchelmann et al., 2017), including grapevine (Konlechner and Sauer, 2016). Albeit one of the main functions of these structures is to protect the plant from, e.g., herbivorous, insects and fungi, the presence of trichomes in the leaf may inhibit the deposition of a homogeneous coating of the matrix. This may lead to an inaccurate ionisation of the ions present on the leaf and consequently a misleading detection/identification of the compounds (Bjarnholt et al., 2014). Having this in mind, and since grapevine leaves possess trichomes which could interfere with the analysis, our first approach was to analyse leaf discs with and without trichomes. Our results showed that leaf discs with no trichomes provide a more accurate analysis of the site-specific compounds. Another important point in sample preparation for MALDI imaging is the matrix. Despite numerous matrices are available to analyse a wide range of biological compounds, it is important to consider the compounds of interest to be analysed in the sample. Selection of an appropriate matrix is a critical point to obtain a high quality MALDI imaging mass spectrum. For metabolomics analysis, the choice of matrices for metabolite analysis is even more complex, since matrix ions often crowd the low-mass range, limiting confident detection of low-molecular weight analyte ions. For the detection of low-weight molecules, α-cyano-4- hydroxycinnamic acid (HCCA), 2,5-dihydroxybenzoic acid (DHB) and 9-aminoacridine (9- AA) have been found to be the better suited (Laugesen and Roepstorff, 2003a; Calvano et al., 2018; Leopold et al., 2018). Besides matrix itself, matrix application, solvent composition (typically methanol or acetonitrile) and mode of application, e.g., sprayer device, movement speed, solvent flow rate, distance between sample and target and/or nozzle temperature, influence the analysis of target molecules from the sample. These parameters are directly related to the size of the matrix crystals formed (Laugesen and Roepstorff, 2003a; Leopold et al., 2018; Schulz et al., 2019). The crystallisation between the matrix and the analyte leads to co-crystals which should be as homogeneous as possible, to assure reproducibility and highest sensitivity. Hence, spray is currently the most widespread matrix application technique. Taking this information into account, in this work, HCCA, DHB and 9-AA were tested with MeOH and ACN as solvents. Our results demonstrate that both solvents can be used for grapevine leaf discs. Also, all matrices appear to be suitable for grapevine leaf disc analysis, within the range of analysis. However, it is important to consider that the intensity of the peaks may be altered according to the matrix used. The obtained molecular mass and the deviation obtained for the ion images of the *m/z* values 365.105 and 381.079 make them identified as a sodiated ([M+Na]^+^) and potasiated ([M+K]^+^), respectively, disaccharide with a molecular formula of C12H22O11, putatively assigned to sucrose. Sucrose is the most common disaccharide in plants and our previous results showed that sucrose increased in *Plasmopara viticola*-infected grapevine leaves 24h post-inoculation (sucrose was identified by FT-ICR-MS with a deviation between 0.2 and 1 ppm as differentially accumulated in inoculated leaves and was further quantified in the homogenised leaves) (Nascimento et al., 2019). Therefore, we can attribute the reconstructed ion images of the *m/z* values 365.105 and 381.079 to sucrose. In this study, the *m/z* values were commonly identified in all of the matrices and solvents tested presenting high levels of abundance in grapevine leaf discs infected with *P. viticola* when comparing with mock inoculated leaf discs.

*Plasmopara viticola* infection occurs when the zoospores germinate and penetrate the stomatal cavity on the lower side of the leaf, forming a substomatal vesicle. This vesicle gives rise to the primary hyphae and mycelium, which grows through intercellular spaces, enclosed by the veins of the leaf (Fröbel and Zyprian, 2019). The haustorium penetrated the parenchyma cell walls, allowing the contact between the pathogen (*P. viticola)* and the host (grapevine leaves) (Gessler et al., 2011; Yin et al., 2017). Also, this structure may function as a site of molecular exchange of effectors and nutrients between the grapevine and *P. viticola* and block grapevine defence signalling pathways (Yin et al., 2017). In fact, our results suggest a link between putatively identified sucrose and the development of *P. viticola* infection structures due to its higher intensity around the veins. Indeed, in grapevine, sucrose is the main sugar-transport molecule, exported from leaves in the phloem (Walker et al., 2021) and there are different hypothesis regarding the roles of sugar transporters in pathogen defence (Bezrutczyk et al., 2018). For example, on *Vitis vinifera* L. ‘Modra frankinja’ leaves infected with Flavescence Dorée phytoplasma, it was observed an increase of sucrose concentration, as well as a higher activity of sucrose synthase (Prezelj et al., 2016). In this interaction, phloem transport is inhibited, resulting in the accumulation of carbohydrates and secondary metabolites that cause a source-sink transition (Prezelj et al., 2016). Also in grapevine, there is an accumulation of sucrose in leaves infected with phytoplasma (Hren et al., 2009).

Up to our knowledge, our results are the first evidence of a metabolite putatively identified as sucrose near to *P. viticola* infection site areas, which lead us to hypothesise that the pathogen, through mycelium development, has gained access to this plant resource, and it is using it to complete its lifecycle (Aked and Hall, 1993; Sutton et al., 1999; Gebauer et al., 2017; Bezrutczyk et al., 2018). This accumulation is in accordance with our previously results on leaf metabolite profiling by NMR, where we reported higher accumulation of sucrose in *V. vinifera* ‘Trincadeira’ in the first 48 hours after inoculation with *P. viticola* (Ali et al., 2012). Also, in the all-leaf tissue analysis of *Vitis vinifera* L. ‘Marselan’, an increase of soluble sugars after 7 dpi (days post-inoculation) was reported (Gamm et al., 2011). The specific location of sucrose needs to be better understood according to the involved biosynthesis pathways and their isomeric composition. Also, sucrose localisation in different *Vitis* genotypes, with different resistance degrees towards pathogens, should be further investigated to understand its role in grapevine defence against pathogens.

The results presented in this article clearly demonstrate the potential of MALDI-FT-ICR-MSI to gain more knowledge on the grapevine-*P.viticola* interaction particularly through the localisation and visual tracking of specific compounds in leaf tissue. Furthermore, our MSI results demonstrate that the analysis methodology works for complex samples, such as grapevine leaves, showing a great potential for application to other plant leaves.

## 5. Conclusion

Grapevine leaves are quite challenging in obtaining high quality MALDI mass spectra images, as leaves are not entirely flat and possess trichomes, influencing the detection of ions. In this work, we have shown that higher ion signal is achieved in leaf discs without trichomes. Three different matrices were selected (HCCA, DHB and 9-AA), whether dissolved in MeOH or ACN, for leaf discs coating and analysis. HCCA showed the highest signal intensity, independently of the solvent.

Reconstructed images by MS allow us to identify sucrose in all the conditions tested.

Sucrose ions were more abundant in the infected leaf discs, comparing to control discs. These results are in accordance with previous studies, which describe a variation of accumulation of sucrose in infected *P. viticola* plants. Up to our knowledge, our results are the first visual evidence of the localisation of sucrose translocation in grapevine leaves linked to the development of *P. viticola* infection structures.

## 6. Conflict of Interest

The authors declare that the research was conducted in the absence of any commercial or financial relationships that could be construed as a potential conflict of interest.

## 7. Author contributions

M.M., A.M., E.P., A.F. and M.S.S. conceived the study and performed the experimental design. M.M., J.C. and J.E.E.D. collected the plant material. M.M. and A.M. per-formed the MALDI MSI analysis. M.M., A.M. and C.M. performed the microscopic analysis. M.M., A.M., A.F. and M.S.S. wrote the manuscript. All authors reviewed the manuscript.

## 8. Funding

The authors acknowledge the support from Fundação para a Ciência e a Tecnologia (Portugal) though the projects PTDC/BAA-MOL/28675/2017, UID/MULTI/04046/2019, UIDB/04292/2020, Investigator FCT programs IF 00819/2015 to Andreia Figueiredo and CEECIND 02246/2017 to Marta Sousa Silva and the PhD grant SFRH/BD/116900/2016 to Marisa Maia. Andréa McCann thank the Excellence of Science Program of the FNRS F.R.S (Rhizoclip EOS2018000802) for financial support. Cedric Malherbe acknowledge support from the F.R.S.-FNRS as Research Associate fellowship. The authors also acknowledge the support from the Portuguese Mass Spectrometry Network (LISBOA-01-0145-FEDER-022125) and the Project EU_FT-ICR_MS, funded by the Europe and Union’s Horizon 2020 research and innovation program under grant agreement nr.731077 (under a staff exchange program). The MALDI FT-ICR SolariX XR instrument was funded by FEDER BIOMED HUB Technology Support (number 2.2.1/996) and the SunChrom sprayer was founded by the European Union’s Horizon 2020 program (EURLipids Interreg Eurogio Meuse-Rhine project supported by the European Regional Development Fund (FEDER)).

## 9. Acknowledgments

The authors acknowledge Gonçalo Laureano for the help in image editing and Prof. Dr. Cristina Figueiredo for the help to operate the lyophilizer.

## Notes

### Competing Interest Statement

The authors have declared no competing interest.

