## Supplementary Material for "Grapevine leaf MALDI-MS imaging reveals the localisation of a putatively identified sucrose metabolite associated to *Plasmopara viticola* development"

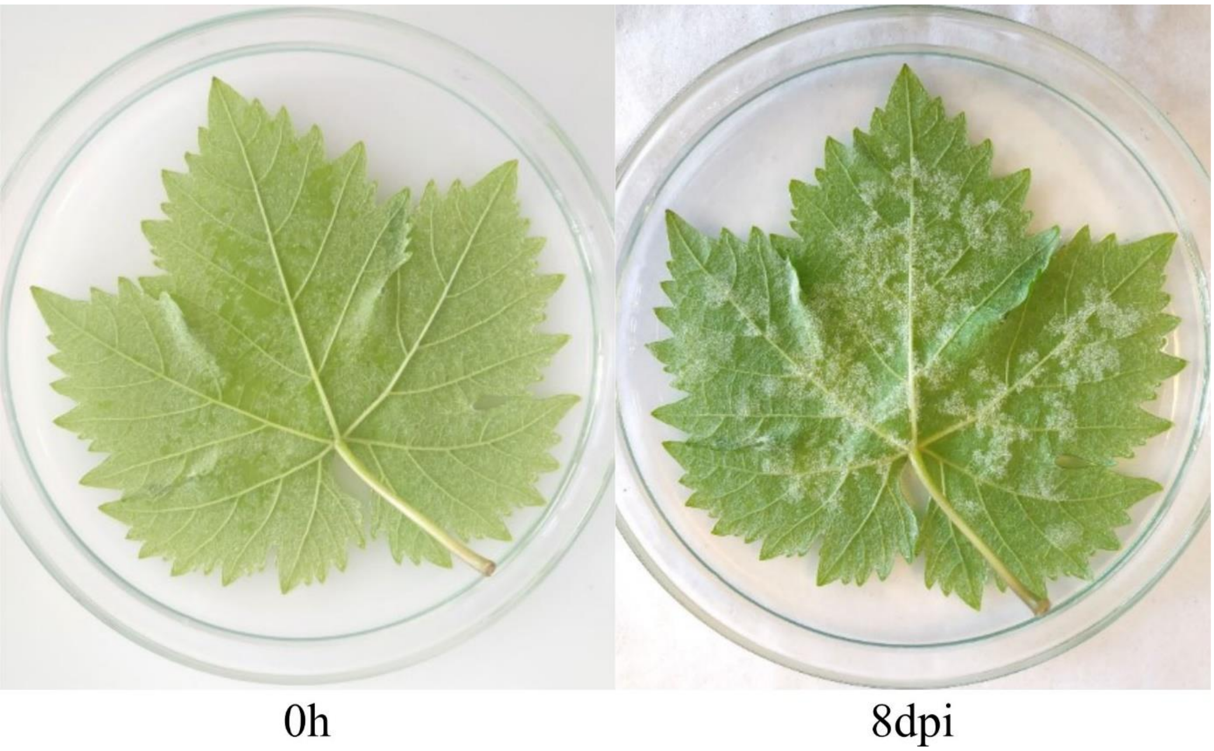


**Supplementary Figure S1.** *V. vinifera* cv Trincadeira phenotype leaves of non-infected (0 h) and 8 days after inoculation (8 dpi) with *P. viticola*.

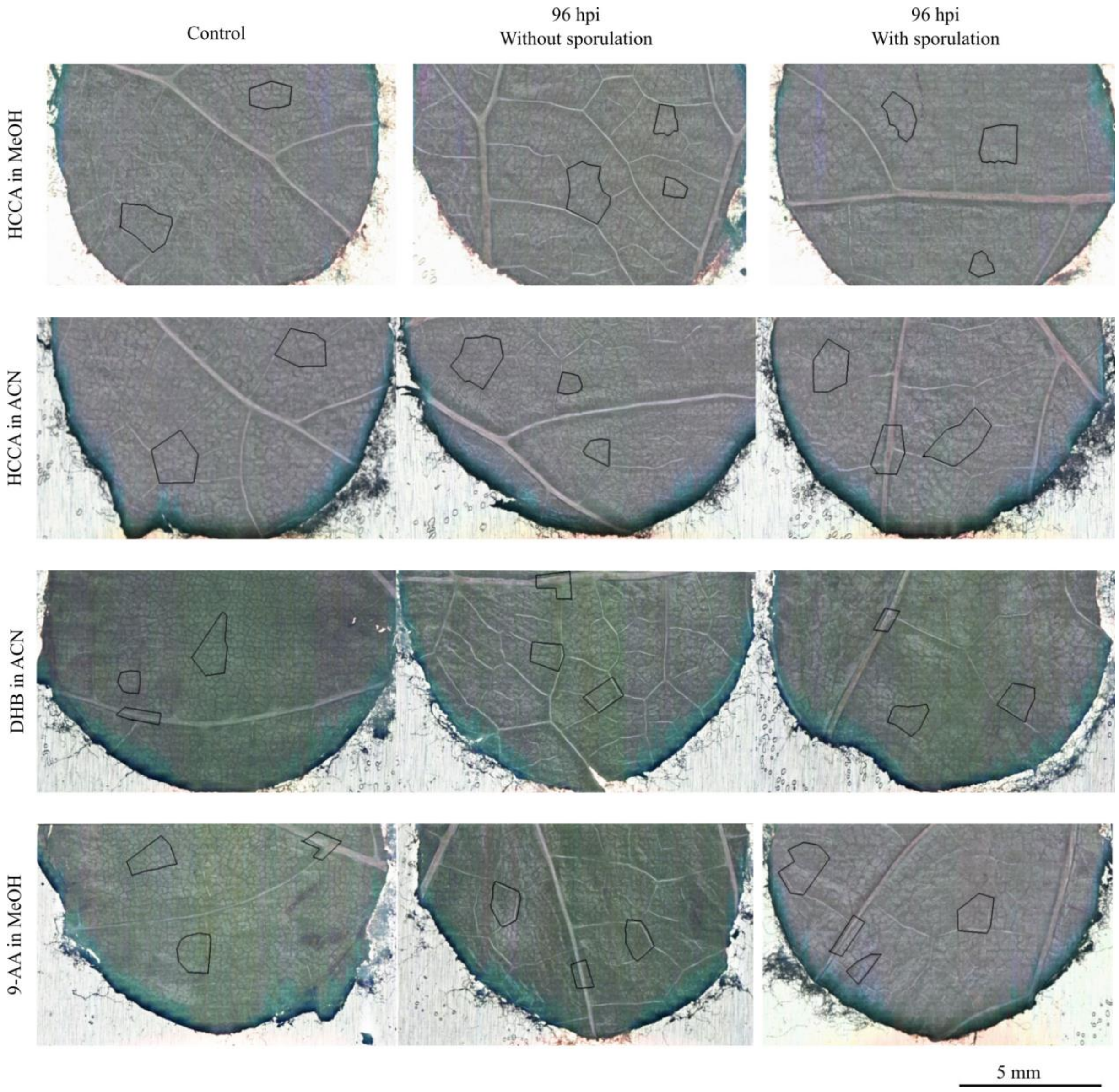


**Supplementary Figure S2.** Microscope images (10 x) of *Vitis vinifera* ‘Trincadeira’. Areas marked with a black line were selected for MALDI-FT-ICR-MS analysis with different matrices and solvents.

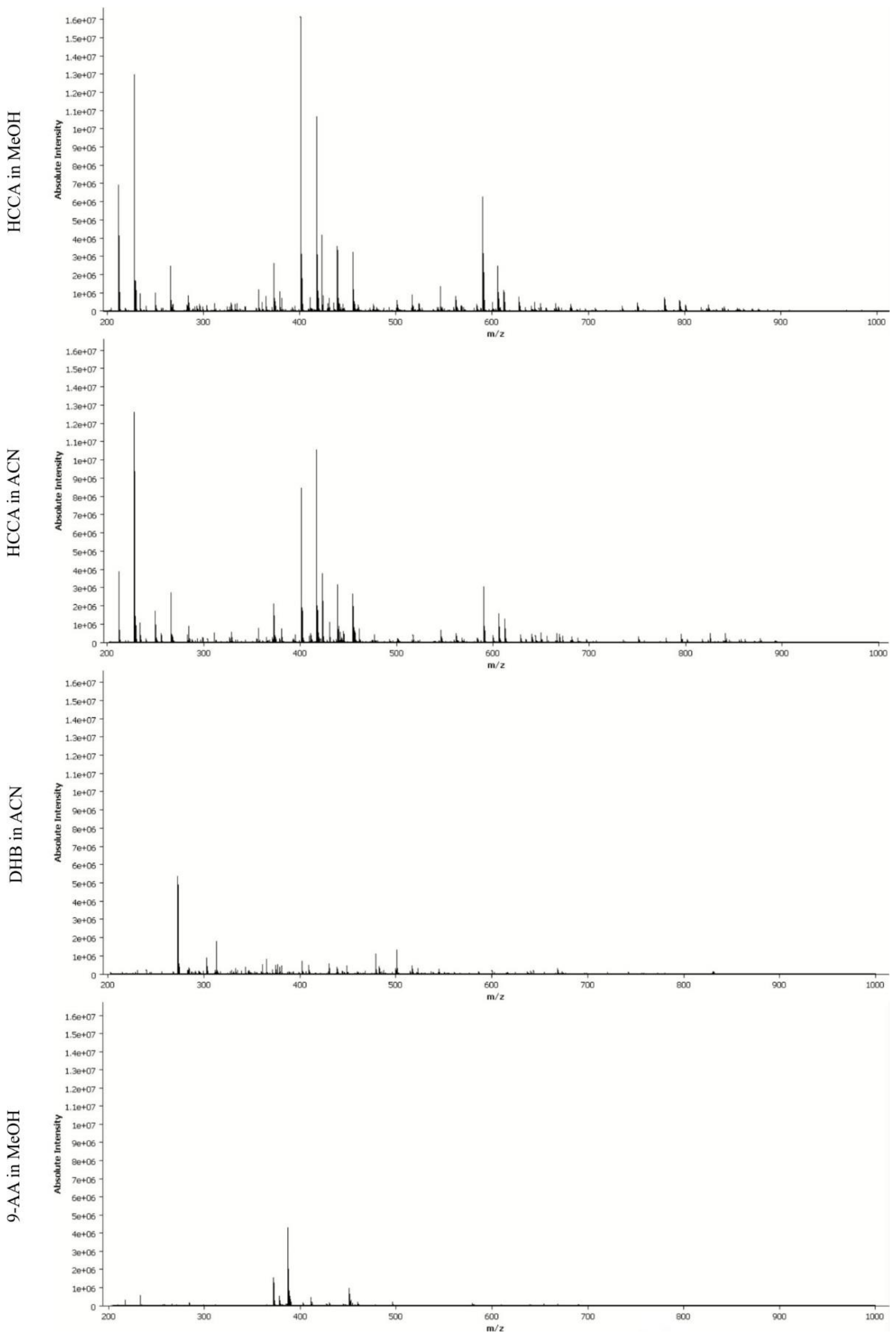


**Supplementary Figure S3.** MALDI-FT-ICR-MS mean mass spectra of *Vitis vinifera* 'Trincadeira' leaf discs (control, 96 hpi without visible *P. viticola* sporulation and 96 hpi with visible sporulation) in positive ionisation mode. Data was normalised by the total ion count.

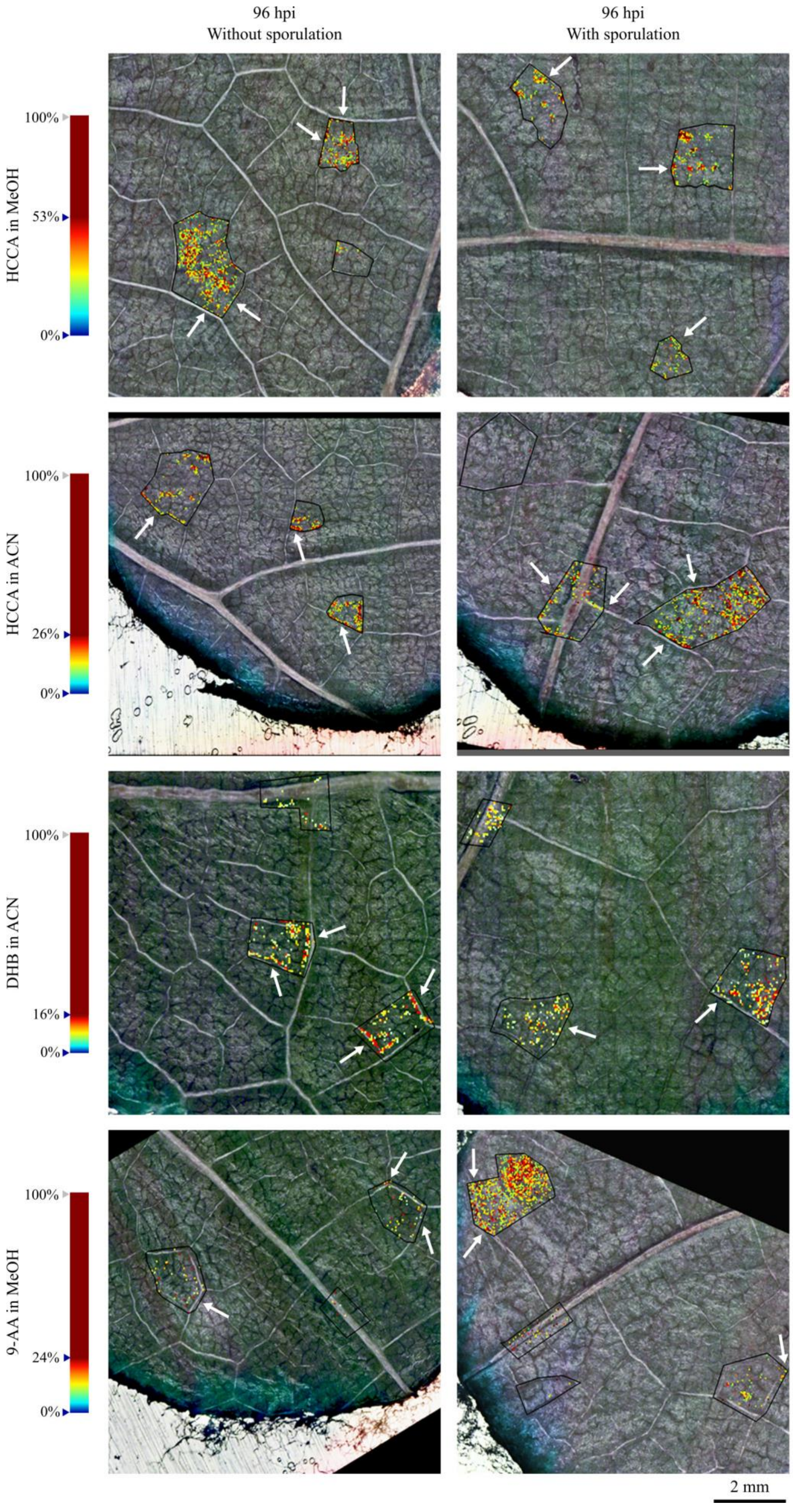
96 hpi
Without sporulation
96 hpi
With sporulation
HCCA in MeOH
100%
53%
0%
HCCA in ACN
100%
26%
0%
DHB in ACN
100%
16%
0%
9-AA in MeOH
100%
24%
0%
2 mm

**Supplementary Figure S4.** Reconstructed ion images of putatively identified sucrose (m/z 365.105, $[M+Na]^+$) in 96 hpi without visible *P. viticola* sporulation and 96 hpi with visible *P. viticola* sporulation. Leaf disc areas detected via MALDI–FT-ICR-MS imaging using HCCA matrix with MeOH and ACN, DHB matrix and 9-AA matrix. The colour scale indicates the absolute intensity of each pixel (arbitrary units). White arrows indicate the visible distribution of putatively identified sucrose along the leaf veins. Spatial resolution: 30µm.
